# Cortical areas involved in grasping and reaching actions with and without visual information: an ALE meta-analysis of neuroimaging studies

**DOI:** 10.1101/2022.06.01.494343

**Authors:** Samantha Sartin, Mariagrazia Ranzini, Cristina Scarpazza, Simona Monaco

## Abstract

The functional specialization of the ventral stream in Perception and the dorsal stream in Action is the cornerstone of the leading model proposed by Goodale and Milner in 1982. This model is based on neuropsychological evidence and has been a matter of debate for almost three decades, during which the dual-visual stream hypothesis has received much attention, including support and criticisms. The advent of functional magnetic resonance imaging (fMRI) has allowed investigating the brain areas involved in Perception and Action and provided useful data on the functional specialization of the two streams. Research on this topic has been quite prolific, yet very little attempt has been made so far to identify consistent neuroimaging results across the available literature. In particular, no meta-analysis has explored the spatial convergence in the involvement of the two streams in Action. The present meta-analysis (N=53) was designed to reveal the specific neural activations associated with Action (i.e., grasping and reaching movements), and the extent to which visual information affects the involvement of the two streams during motor control. Our results provide a comprehensive view of the consistent and spatially convergent neural correlates of Action based on neuroimaging studies conducted over the past two decades. We discuss our results in light of the well-established dual-visual stream model and frame these findings in the context of recent discoveries obtained with advanced fMRI methods, such as multivoxel pattern analysis.

## 1. Introduction

Over the last three decades, the investigation of the neuroanatomical substrates of hand action has received increasing attention. Indeed, hand actions not only allow us to interact with our surroundings, but also enable us to satisfy our basic needs. A deep and comprehensive understanding of the neural mechanisms underlying hand action is crucial for advancements in many research areas, such as the ever-growing field of brain-computer interfaces for individuals who have limited or no ability to perform volitional movements.

One of the most prominent theories about action and perception was put forward by Goodale and Milner in 1992 and is based on behavioural and neuropsychological findings showing a specialization of the dorsal stream in action and the ventral stream in perception (Goodale & Milner, 1992). Specifically, the original model describes the functional specialization of the parietal cortex in processing spatial information that is relevant for planning and executing action, and the temporal-occipital cortex in recognition of contents. The proposed two-visual streams model has received much interest, including support and criticisms, and it has been a matter of debate for almost three decades (see Freud et al., 2016; Whitwell et al., 2014).

The advent of functional magnetic resonance imaging (fMRI) has allowed investigating the brain areas involved in action and perception and provided useful data on this topic. Despite the challenges related to studying the execution of motor actions with neuroimaging techniques, among which the confined environment of the MR and the risk of inducing motion artifacts, research on the functional specialization of the two streams has been quite prolific (e.g., Cavina-Pratesi et al., 2007; Culham et al., 2003; Króliczak et al., 2007, 2008; Singhal et al., 2013). Yet very little attempt has been made so far to determine the consistency in neuroimaging results across the published studies, with only one recent study which summarizes the existing data with systematic research and a meta-analytical approach (Ranzini et al., 2022).

In addition to the specialization of the dorsal and ventral stream in action and perception, respectively, according to the dual-stream theory both streams are involved in processing vision for action, but with different purposes (Goodale & Milner, 1992; Milner & Goodale, 2008). Specifically, while the dorsal visual stream is specialized in the online control of visually-guided actions, the ventral visual stream processes short-term maintenance of the object representation in memory, and, as a consequence, it is thought to support the guidance of delayed actions in absence of online visual information (i.e. when the brief visual presentation of a stimulus and the action towards it are separated by a delay). Seminal evidence comes from neuropsychological observations of patients suffering from optic ataxia (Perenin & Vighetto, 1988), on the one side, and patients suffering from visual agnosia, on the other side (Riddoch & Humphreys, 1987). Optic ataxia is due to dorsal stream lesions and consists of a deficit in reaching towards objects in the visual periphery. Visual agnosia is caused by ventral stream lesions which impair the visual recognition of objects and shapes. Importantly, while optic ataxia patients perform more accurate actions with than without a delay (Milner et al., 1999, 2001), visual agnosia patients show the opposite pattern (Goodale et al., 1994), in line with the idea that while the dorsal stream is crucial for online control of immediate actions, the ventral stream permits maintenance of object representation in memory for delayed actions.

Several studies using different techniques, such as fMRI and transcranial magnetic stimulation (TMS), supported and extended the dual-stream theory by showing that while dorsal stream areas play a role in immediate and delayed actions, the ventral stream might have a more prominent role in delayed actions only. In particular, neuroimaging studies showed that during delayed actions ventral and dorsal stream areas are re-activated when a movement is performed in absence of visual information (Himmelbach et al., 2009; Monaco et al., 2017; Singhal et al., 2013). Further, TMS studies determined that while the dorsal stream has a causal role in performing immediate and delayed actions (Cohen et al., 2009; Smyrnis et al., 2003), the ventral stream is crucial for delayed but not immediate actions (Cohen et al., 2009). In addition to these findings, some fMRI studies showed that the early visual cortex is reactivated during delayed actions in the dark (Chen et al., 2014; Monaco et al., 2017; Singhal et al., 2013).

One of the controversies about previous findings on the involvement of dorsal and ventral stream areas in hand actions arises from the fact that not all results point to the involvement of the ventral stream and the early visual cortex in delayed actions without online visual feedback. Indeed, some fMRI studies found reactivation in dorsal but not ventral stream areas during the execution of an action after a delay (Fiehler et al., 2011), while other studies found reactivation in the ventral stream and EVC during action execution in the dark after a delay (Chen et al., 2014; Monaco et al., 2017; Singhal et al., 2013). Further, the reactivation in visual areas for delayed actions in the dark was higher for grasping than reaching actions, perhaps because grasping requires the retrieval of more detailed information, such as size and shape) than reaching. Yet, no attempt has been made to assess the consistency of results across the literature in a systematic manner.

In this study we exploited the potential of coordinate-based Activation Likelihood Estimation (ALE) meta-analysis on neuroimaging studies (Laird et al., 2005; Turkeltaub et al., 2002), to explore the neural bases of hand reaching and grasping, performed with and without online visual feedback (i.e., after a delay following the presentation of the target object or in total darkness). We focused on studies investigating hand reaching and grasping because reach-to-grasp is probably the most representative human skilled-action, and it has been extensively investigated to test and confirm the dual-visual stream theory (Goodale & Milner, 1992). We based our literature search and article selection on a recent study by Ranzini et al. (2022), where a systematic review and meta-analysis of neuroimaging studies on the execution of reach and grasp actions has been provided. Meta-analytical studies have proven to be important for the assessment of consistency across neuroimaging studies (Wager et al., 2009). Indeed, they enable us to get a summary of the brain areas (i.e., activation clusters) that are active during a particular task, or involved in a cognitive domain, across all the studies published so far.

The main objective of the current study was to elucidate the brain areas consistently recruited during the execution and guidance of skilled actions, specifically hand reaching and grasping actions, when visual information of the target object and/or the hand is available, and when it is not. More specifically, the present work aimed at addressing two main aims. First, we investigated whether activation in the ventral stream and early visual cortex is found only during the execution of reaching and grasping actions with online visual feedback, as would be expected because of the presence of a visual input, or also when delayed actions are executed in complete darkness after a brief presentation of a stimulus (aim 1). To address this aim, we performed two meta-analyses, one including all the studies investigating the execution of reaching and grasping with online visual feedback, and the other one including the experiments exploring the execution of the same actions without visual feedback, and therefore relying on memory. We then directly compared (through a contrast analysis) the two sets of studies (i.e., reaching and grasping with vision vs. without vision) to identify the visual areas that show consistently higher activation for actions executed with as compared to without visual input. Given the well-known role of the ventral stream and early visual cortex in visual perception, we expected to find it consistently activated during actions executed with online visual feedback, as the visual input of the target or the hand approaching the target are being processed (Bracci et al., 2010; Malach et al., 1995). The critical question was whether the ventral stream and early visual areas also show consistent activation during actions performed without visual feedback, given the divergent results on the involvement of ventral stream areas in delayed actions (i.e., Fiehler et al., 2011; Singhal et al., 2013). Second, we assessed whether reaching and grasping differentially recruit ventral and dorsal stream areas (aim 2). Some studies have shown that grasping elicits higher activation than reaching actions in the ventral stream and early visual cortex (Monaco et al., 2017; Singhal et al., 2013). This might be related to the fact that grasping, but not reaching, requires detailed information about the object properties, such as size and shape, to perform an accurate movement. As such, we hypothesized consistently higher activation in ventral stream and early visual areas for grasp than reach tasks. To address this aim, we performed two meta-analyses including the experiments investigating: grasping, on one side, and reaching, on the other side. Lastly, we ran a contrast analysis between grasping and reaching (i.e., grasping vs. reaching).

## 2. Materials and methods

### 2.1 Studies selection

The procedure used for the selection of the studies is described in detail in Ranzini et al. (2022). The data associated with this study is publicly available and can be found at: https://osf.io/48w69/?view_only=633d3ad74a1346ab86b65d6766f79753.

To summarize, the literature search was conducted until December 30, 2020 (for a detailed description of the literature searching process, see PRISMA flow diagram, Annex A). To the best of our knowledge, there was no study published after this date that could be included in this meta-analysis. We identified 565 studies through a database search with Pubmed and bioRxiv on hand reaching and grasping. Further, 454 studies were identified with the use of the “related articles” function, available in the Pubmed database, and the backward and forward snowballing search strategy, i.e., reference list and citations of primary articles, reviews and meta-analyses. This selection process led to a total of 1020 studies. After removing duplicates, a total of 954 studies were originally identified to undergo further scrutiny at a later stage.

Studies had to respect the following inclusion criteria to be included in the current meta-analysis:

- to have written the paper in the English language.
- to use a hand reaching and/or grasping task.
- to investigate brain activity during the action execution phase (i.e., studies that focused on brain activity during the planning phase were excluded). This ensured consistency across the selected studies, where the elicited activation reflected somatosensory feedback and motor outputs, which are absent during the planning phase preceding action execution. This criterion was added to the ones used in the study by Ranzini and colleagues (2022), where the execution as well as the planning phase preceding the movement were considered. As such, five studies were excluded from our meta-analysis (Beurze et al., 2007, 2009; Chapman et al., 2007; Chen et al., 2014; Majdandžić et al., 2007).
- to use functional magnetic resonance imaging (fMRI) or positron emission tomography (PET) to collect data about neural activity.
- to have conducted whole-brain analyses (e.g., studies that use a region of interest (ROI) approach were excluded, as it focuses on predefined areas of the image rather than reporting all activated clusters and could thus bias the result of the meta-analysis; Müller et al., 2018).
- to have performed univariate analyses (i.e., papers that conducted multivoxel pattern analysis or functional connectivity analyses were not included).
- to report a contrast of reaching or grasping larger than the one in the control condition.
- to report findings in either Talairach (Talairach & Tournoux, 1988) or Montreal Neurologic Institute (MNI) coordinate space (i.e., studies not reporting results in a standardized coordinate space were excluded).
- to have included only healthy adults in the experiment.
- to test a sample size of at least 5 participants.

### 2.2 Systematic review

The literature was screened in detail and the articles that met the inclusion criteria were selected in accordance with PRISMA guidelines (Moher et al., 2009) by Ranzini and colleagues (2022). We checked that the screening procedure was in line with the updated PRISMA guidelines that have been recently published (Page et al., 2021). In addition, we followed recent recommendations on how to conduct a proper neuroimaging meta-analysis (Müller et al., 2018). The screening procedure is described in more detail in the PRISMA flow diagram available in Annex A. For the current meta-analysis, 53 studies met the inclusion criteria reported in the previous section. The complete list of included studies is presented in Annex B.

Data were extracted from the studies and then checked. We then created a database containing the following information of the selected articles on hand reaching and grasping actions: the sample size, the percentage of females, the mean age of participants, the technique (either fMRI or PET), the experimental task (only grasp, only reach, or reach and grasp), the control task, additional information about the task and stimuli, the relevant contrast selected, the coordinate system, the coordinates of foci and their anatomical labels, the p-value criteria (corrected, uncorrected), and the related statistic (z score, t value).

In the case of multiple contrasts performed in a single study and on the same group of participants, only the most relevant one was considered (i.e., the contrast that best represents the process under investigation in the present meta-analysis). This approach prevents from creating dependence across experiment maps that negatively impacts the validity of meta-analytic results (Müller et al., 2018). As a result of the application of this approach, we eventually selected only one contrast from each of the eligible studies, thus yielding a final list of 53 experiments (i.e., contrasts), consisting of 528 foci, included in the current meta-analysis. We then divided the studies into four categories consisting of two movements (Grasp and Reach) and two levels of visual information (Vision and No vision), as described here below.

### 2.3 Data categorization

For the purpose of the current meta-analysis, we further analysed each of the 53 studies in order to extract additional information about the experimental task and the contrast used.

Reaching tasks consisted of moving the hand towards the object with the pointing finger or the knuckles. Grasping tasks consisted of moving the hand towards the object and grasping it with a precision or a whole hand grasp. While some of the studies in this meta-analysis employed either reach or grasp tasks, others included both movement types. Some studies also included ad-hoc control conditions (i.e., passive viewing of the target object). Tasks consisting in moving the arm and hand to the target, without a grip component, were categorized as Reach. Tasks that included the grip component, with or without the movement of the arm towards the target, were categorized as Grasp. For example, the contrasts of: (Grasp vs. Reach), and (Grasp vs. Passive viewing) were both labelled as Grasp, as they both included the grip component (note that this procedure partially differs from the one of Ranzini et al. (2022), where a distinction between reach-only, grasp-only, and reach-to-grasp studies was made).

We then assessed whether the experimental paradigm required participants to perform hand actions in total darkness (i.e., participants could not see their moving hand or the target object; No vision condition) or in a dimly light room (i.e., participants could see their own hand moving and the target object or a visual stimulus projected onto a screen; Vision condition). For experiments performed under dim light illumination sufficient to process visual stimuli during action execution, we determined whether the contrast used in the study allowed subtracting neural activity elicited by the visual stimuli and/or the hand performing the movement. If so, we included these contrasts in the No vision condition along with the studies in which participants performed hand actions in complete darkness.

As a result, we classified the selected studies into the following categories: 1) reaching experiment in which neural activity elicited by the vision of the moving hand and/or the target object during action execution was not present (i.e., actions were performed in complete darkness) or was subtracted by the contrast (Reach No vision; number of studies (N)=13); 2) reaching experiment in which visual information about the hand or the target was present (Reach Vision; N=3); 3) grasping experiment in which neural activity elicited by the visual information of the moving hand and/or the target object was not present (i.e., actions were performed in complete darkness) or was subtracted by the contrast (Grasp No vision; N=20); 4) grasping experiment in which vision was present (Grasp Vision; N=17).

### 2.4 Activation likelihood estimation meta-analysis

We conducted a coordinate-based meta-analysis (CBMA) which uses the coordinates of activation peaks (i.e., activation foci) reported in a standardized coordinate space. We employed the ALE method (Laird et al., 2005; Turkeltaub et al., 2002) to conduct the coordinate-based meta-analysis of selected fMRI and PET experiments. In particular, the revised version of the ALE algorithm (Eickhoff et al., 2009, 2017) was run with BrainMap GingerALE software version 3.0.2 (Research Imaging Institute; http://brainmap.org/ale/). The MNI coordinates of activation peaks were converted into Talairach space before performing the meta-analysis.

The ALE algorithm aims at evaluating the brain areas of spatial convergence of activated foci across neuroimaging experiments using the coordinates of the peak activations extracted from individual studies. In particular, the algorithm models the reported activated foci of each experiment as three-dimensional Gaussian probability distributions. The number of participants in each experiment is considered to compute the size of the probability distributions. The uncertainty associated with the spatial location of activated foci due to between-study variances (e.g., between-subject and between-template variances; Eickhoff et al., 2009) is considered, quantified and used by the ALE algorithm to compute the width of each Gaussian distribution. The probability distributions of all activation foci extracted from an experiment are then combined voxelwise to obtain a Modelled Activation (MA) map, that is a map (i.e., 3D volume) of activation likelihood that is generated for each included experiment. For each meta-analysis all the MA maps are combined voxelwise to create an ALE map. Each voxel of this image contains an ALE score which represents the spatial convergence of activated foci at exactly that position (Eickhoff et al., 2009). The ALE scores are then tested against a null hypothesis according to which the concordance in spatial activation between experiments can occur by chance and is therefore random (noise; Eickhoff et al., 2016), by applying a random-effects spatial inference (i.e., random effects model) instead of a fixed-effects inference to evaluate the agreement on activation peaks across studies. The ALE algorithm uses a permutation procedure to assign each voxel a P value which stems from the probabilities of obtaining an ALE value not equal to the ALE value of the very same voxel based on the null-distribution. We employed Mango software (http://ric.uthscsa.edu/mango/), a multi-image analysis program, to visualize the results of the meta-analysis by overlaying ALE maps onto an anatomical image in Talairach space.

#### 2.4.1 Single dataset and contrast analyses

We ran four single dataset and two contrasts analyses to examine the areas involved in the execution of reaching and grasping movements with and without the availability of visual information. In order to examine which areas are consistently involved in visual and non-visual reaching and grasping actions, we performed two meta-analyses separately for each of the two visual conditions across action types: 1) Reach and Grasp Vision, and 2) Reach and Grasp No vision. Further, to investigate which areas are selectively involved in online visual processing during action execution, we performed a contrast analysis of: 3) Reach and Grasp Vision > Reach and Grasp No vision. To investigate the cortical areas specifically involved in grasping and reaching tasks, regardless of the availability of visual information, we ran two single dataset analyses for each action type (Grasp, Reach) across the two visual conditions: 4) Grasp Vision and No vision, 5) Reach Vision and No vision. Lastly, to determine which areas consistently show higher activation for grasping than reaching movements, we ran the contrast analysis of: 6) Grasp > Reach.

For the single dataset meta-analyses, all the resulting statistical ALE maps were thresholded by means of a cluster-level family-wise error (cFWE) correction at p < 0.05 (5,000 permutations) with a cluster-forming threshold of p < 0.001 (uncorrected), in line with the latest recommendations for neuroimaging meta-analyses (Müller et al., 2018).

To perform the contrast analyses, we followed the recommendation of Eickhoff et al. (2016) according to which two datasets are comparable when one is at most four times bigger than the other one (and vice versa), in terms of number of studies. Therefore, the sample size of studies included in the contrast analyses was: Reach and grasp with vision (N=20) vs. Reach and grasp without vision (N=33), and Grasp (N=37) vs. Reach (N=16). During the analysis, the ALE algorithm repeatedly and randomly splits the pooled list of foci into two separate sets of data while keeping their original sizes. Afterwards, an ALE map is generated for each new dataset, and one is subtracted from the other one (and vice versa); eventually, for each voxel the difference is computed between this new experimental ALE map and the original ALE map. For the current meta-analysis, the ALE algorithm used 10,000 permutations to perform the contrast-analyses. The uncorrected threshold and the minimum cluster volume were set at p < 0.05 and to 100 mm^3, respectively.

## 3. Results

An overview of all areas involved in the execution of reaching and grasping actions, regardless of whether or not vision is available, can be found in Ranzini et al. (2022) (Hand Reaching and Grasping: Figure 2, Panel b; Hand Grasping: Figure 4, Panel a; Hand Reaching, Figure 4, Panel b).

**Figure 1.**
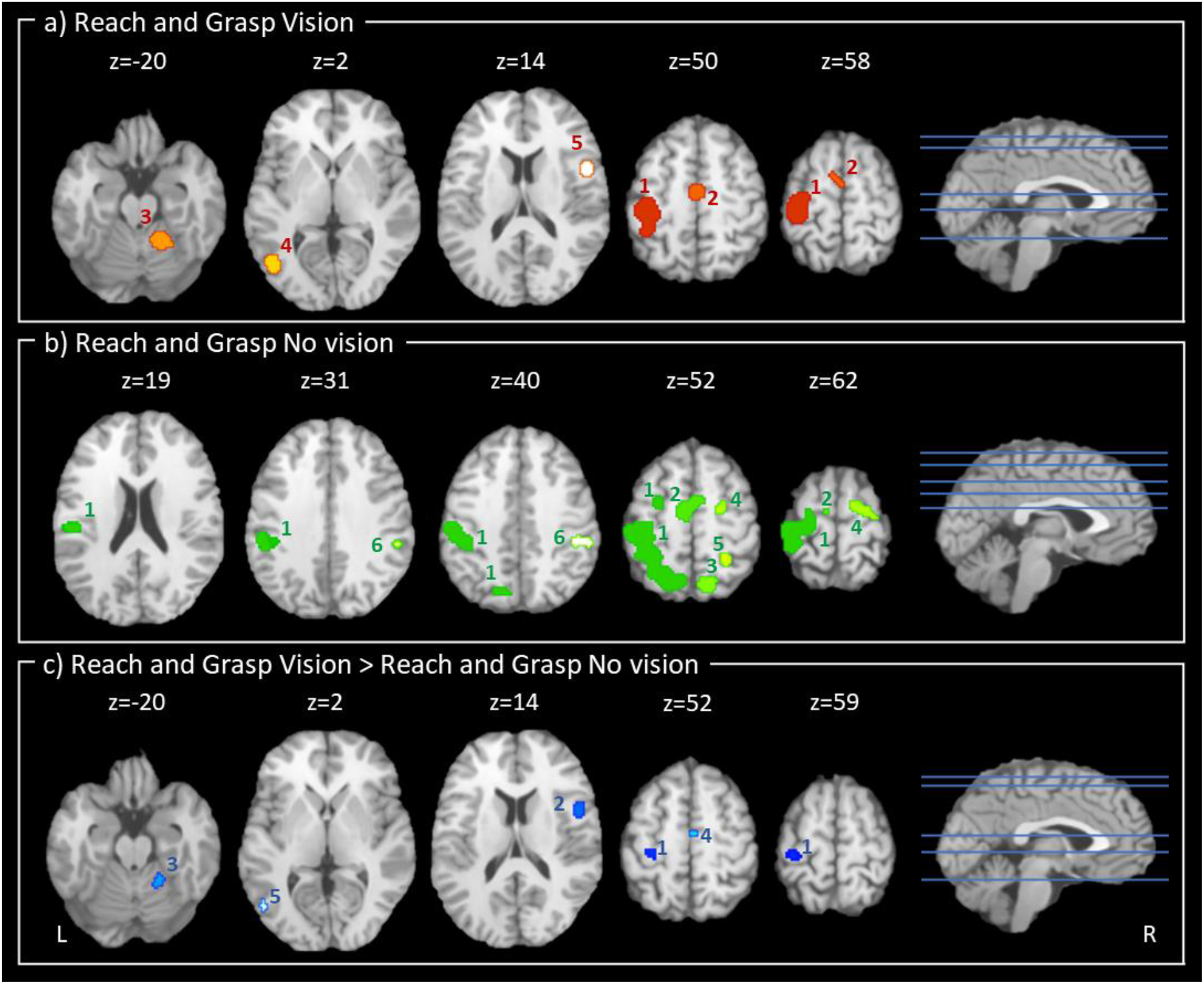
Schematic representation of the results for the Reach and Grasp Vision meta-analysis (Panel a), Reach and Grasp No vision meta-analysis (Panel b), and the Reach and Grasp Vision and Reach and Grasp No vision contrast analysis (i.e., Reach and Grasp Vision > Reach and Grasp No vision; Panel c). Results are shown in the axial view. TAL z coordinates are shown above the slices. Numbers within the slices (1-6) refer to clusters (Panel a: 1=left precentral gyrus, left postcentral gyrus, left inferior parietal lobule, 2=left and right medial frontal gyrus, 3=right cerebellum, 4=left middle temporal gyrus, left inferior temporal gyrus, left middle occipital gyrus, 5=right precentral gyrus, right inferior frontal gyrus; Panel b: 1=left insula, left postcentral gyrus, left inferior parietal lobule, left supramarginal gyrus, left precuneus, left precentral gyrus, left superior parietal lobule, left sub-gyral, left middle frontal gyrus, left superior frontal gyrus, 2=left and right medial frontal gyrus, 3=right precuneus, right superior parietal lobule, 4=right sub-gyral, right middle frontal gyrus, right superior frontal gyrus, right precentral gyrus, 5=right precuneus, right sub-gyral, 6=right inferior parietal lobule, right supramarginal gyrus; Panel c: 1=left precentral gyrus, left postcentral gyrus, 2=right precentral gyrus, right inferior frontal gyrus, right insula, 3=right cerebellum, 4=right medial frontal gyrus, 5=left inferior temporal gyrus, left middle occipital gyrus).

**Figure 2.**
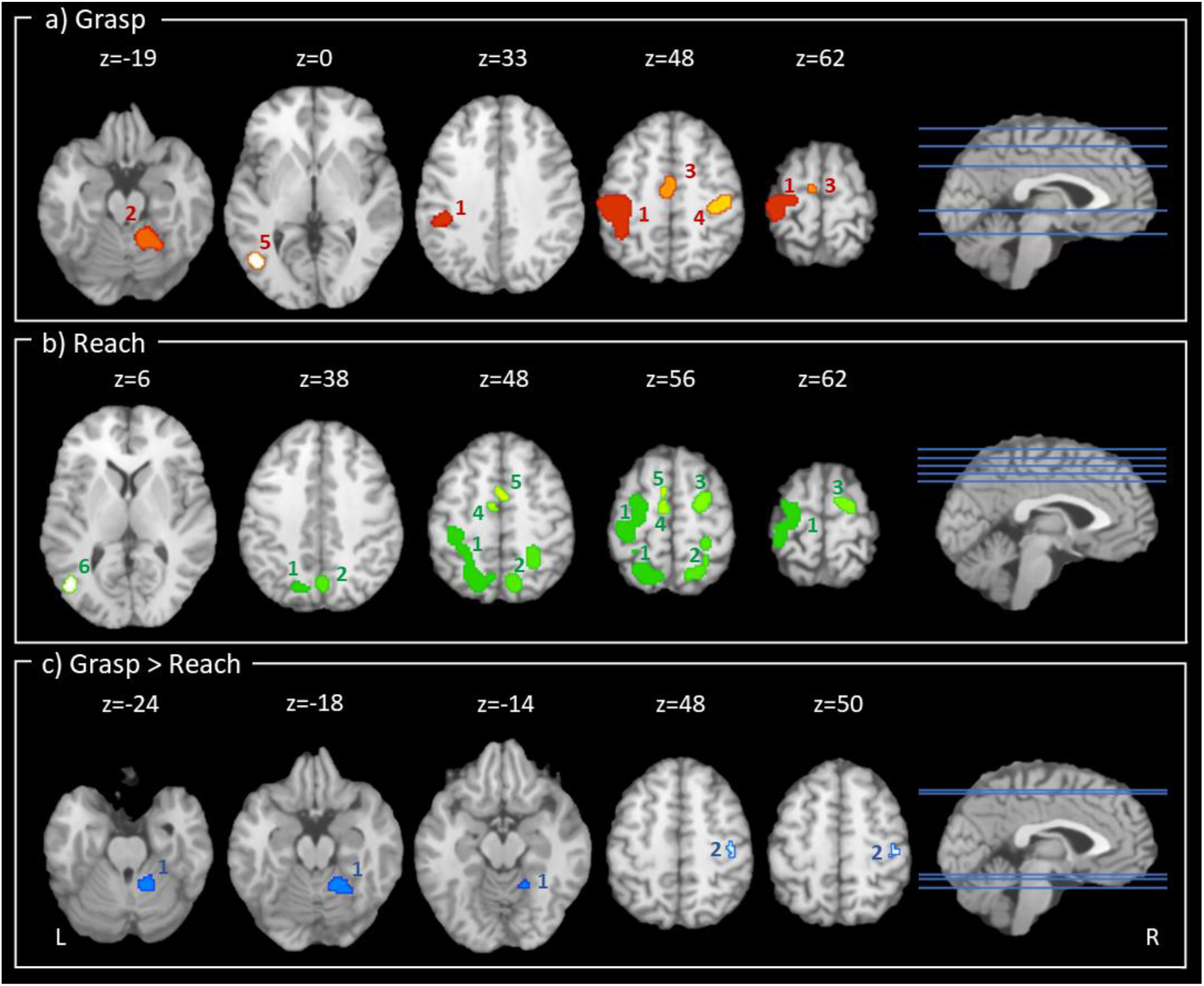
Schematic representation of the results for the Grasp meta-analysis (Panel a), the Reach meta-analysis (Panel b), and the Grasp and Reach contrast analysis (i.e., Grasp > Reach; Panel c). Results are shown in the axial view. TAL z coordinates are shown above the slices. Numbers within the slices (1-6) refer to clusters (Panel a: 1=left precentral gyrus, left postcentral gyrus, left inferior parietal lobule, left supramarginal gyrus, 2=right cerebellum, 3=left and right medial frontal gyrus, left and right paracentral lobule, 4=right precentral gyrus, right postcentral gyrus, 5=left Inferior temporal gyrus, left inferior occipital gyrus, left middle occipital gyrus, left middle temporal gyrus, 6=right precentral gyrus, right inferior frontal gyrus; Panel b: 1=left precuneus, left inferior and superior parietal lobule, left precentral and postcentral gyrus, left sub-gyral, left paracentral lobule, left middle frontal gyrus, left superior frontal gyrus, 2=left and right precuneus, right superior and inferior parietal lobule, right sub-gyral, right paracentral lobule, right postcentral gyrus, 3=right middle frontal gyrus, right sub-gyral, right superior frontal gyrus, right middle frontal gyrus, right precentral gyrus 4=left medial frontal gyrus, 5=left and right cingulate gyrus, left and right medial frontal gyrus, 6=left middle occipital gyrus, left middle temporal gyrus; Panel c: 1=right cerebellum, 2=right postcentral gyrus).

### 3.1 Meta-analytical map of Reach and Grasp Vision

The Reach and Grasp Vision ALE meta-analysis included a total of 298 subjects, and 168 foci extracted from 20 eligible experiments. Results showed five significant clusters (Figure 1, Panel a; Table 1).

**Table 1.** Results of the single dataset meta-analysis on Reach and Grasp Vision. TAL: Talairach; Hemi: hemisphere; BA: Brodmann Area; Cluster size: size of clusters in mm^3.

| Cluster | Cluster size | ALE value | p value | z-score | Hemi | Anatomical Labelling | BA | TAL coordinates |  |  |
| --- | --- | --- | --- | --- | --- | --- | --- | --- | --- | --- |
|  |  |  |  |  |  |  |  | x | y | z |
| 1 | 5,776 | 0.048 | <0.001 | 8.81 | L | Precentral gyrus | 4 | -32 | -30 | 56 |
|  |  | 0.017 | <0.001 | 4.36 | L | Inferior parietal lobule | 40 | -38 | -42 | 52 |
| 2 | 1,280 | 0.023 | <0.001 | 5.35 | R | Medial frontal gyrus | 6 | 2 | -12 | 50 |
|  |  | 0.012 | <0.001 | 3.48 | L | Medial frontal gyrus | 6 | -6 | 0 | 58 |
| 3 | 1,264 | 0.023 | <0.001 | 5.37 | R | Cerebellum (anterior lobe, dentate) | - | 16 | -50 | -20 |
| 4 | 1,064 | 0.022 | <0.001 | 5.32 | L | Inferior temporal gyrus | 37 | -44 | -68 | 2 |
| 5 | 720 | 0.019 | <0.001 | 4.78 | R | Inferior frontal gyrus | 44 | 50 | 6 | 14 |

The most significant peaks of activity were located in the left precentral gyrus (cluster 1; TAL coordinates: -32, -30, 56, BA4), the right medial frontal gyrus (cluster 2; TAL coordinates: 2, -12, 50, BA6), the right cerebellar dentate (cluster 3; TAL coordinates: 16, -50, -20), the left inferior temporal gyrus (cluster 4; TAL coordinates: -44, -68, 2, BA37), and the right inferior frontal gyrus (cluster 5; TAL coordinates: 50, 6, 14, BA44). Cluster 1 (5,776 mm^3) showed two activation peaks in the left hemisphere, and it extended from the postcentral gyrus (42.3% of experiments) to the precentral gyrus (38.4%), and the inferior parietal lobule (19.4%). Cluster 2 (1,280 mm^3) consisted of two activation peaks, and it spanned both the left and right hemispheres (52% and 48% of experiments, respectively); more precisely, it was located in the medial frontal gyrus (BA6; 97.3%), and it spread slightly to the paracentral lobule (BA31; 2.7%). Cluster 3 (1,264 mm^3) was found with one activation peak in the right cerebellar hemisphere; the cluster spanned the anterior lobe (98.7%), and slightly spread to the posterior lobe of the cerebellum (1.3%). Cluster 4 (1,064 mm^3) consisted of one activation peak in the left hemisphere; it was primarily located in the inferior temporal gyrus (40.4%), the middle temporal gyrus (29.8%), and the middle occipital gyrus (29.8%). Cluster 5 (720 mm^3) was found with one activation peak in the right hemisphere; it was located mainly in the precentral gyrus (BA44; 66.7%), and the inferior frontal gyrus (BA6; 33.3%).

### 3.2 Meta-analytical map of Reach and Grasp No vision

The Reach and Grasp No vision ALE meta-analysis included a total of 476 subjects, and 360 foci extracted from 33 eligible experiments. Results showed six significant clusters (Figure 1, Panel b; Table 2).

**Table 2.** Results of the single dataset meta-analysis on Reach and Grasp No vision. TAL: Talairach; Hemi: hemisphere; BA: Brodmann Area; Cluster size: size of clusters in mm^3.

| Cluster | Cluster size | ALE value | p value | z-score | Hemi | Anatomical Labelling | BA | TAL coordinates |  |  |
| --- | --- | --- | --- | --- | --- | --- | --- | --- | --- | --- |
|  |  |  |  |  |  |  |  | x | y | z |
| 1 | 21,288 | 0.034 | <0.001 | 6.25 | L | Postcentral gyrus | 3 | -34 | -30 | 50 |
|  |  | 0.033 | <0.001 | 6.08 | L | Postcentral gyrus | 3 | -38 | -26 | 54 |
|  |  | 0.030 | <0.001 | 5.75 | L | Precuneus | 7 | -20 | -62 | 52 |
|  |  | 0.029 | <0.001 | 5.59 | L | Postcentral gyrus | 2 | -46 | -26 | 48 |
|  |  | 0.029 | <0.001 | 5.55 | L | Postcentral gyrus | 40 | -34 | -34 | 58 |
|  |  | 0.027 | <0.001 | 5.34 | L | Inferior parietal lobule | 40 | -36 | -40 | 56 |
|  |  | 0.024 | <0.001 | 4.91 | L | Superior parietal lobule | 7 | -28 | -58 | 54 |
|  |  | 0.023 | <0.001 | 4.81 | L | Precentral gyrus | 4 | -22 | -24 | 62 |
|  |  | 0.023 | <0.001 | 4.70 | L | Superior parietal lobule | 7 | -10 | -64 | 54 |
|  |  | 0.021 | <0.001 | 4.39 | L | Sub-gyral | 6 | -24 | -6 | 56 |
|  |  | 0.020 | <0.001 | 4.31 | L | Inferior parietal lobule | 40 | -42 | -34 | 36 |
|  |  | 0.019 | <0.001 | 4.21 | L | Postcentral gyrus | 40 | -52 | -24 | 22 |
| 2 | 3,536 | 0.029 | <0.001 | 5.65 | L | Medial frontal gyrus | 6 | -6 | -12 | 54 |
|  |  | 0.020 | <0.001 | 4.33 | R | Medial frontal gyrus | 6 | 6 | -2 | 48 |
|  |  | 0.020 | <0.001 | 4.24 | R | Medial frontal gyrus | 6 | 2 | -4 | 48 |
| 3 | 2,056 | 0.030 | <0.001 | 5.80 | R | Precuneus | 7 | 12 | -68 | 48 |
|  |  | 0.015 | <0.001 | 3.47 | R | Superior parietal lobule | 7 | 22 | -60 | 56 |
| 4 | 1,744 | 0.019 | <0.001 | 4.14 | R | Middle frontal gyrus | 6 | 22 | -8 | 56 |
|  |  | 0.018 | <0.001 | 4.00 | R | Superior frontal gyrus | 6 | 16 | -8 | 62 |
|  |  | 0.016 | <0.001 | 3.56 | R | Precentral gyrus | 6 | 30 | -16 | 60 |
| 5 | 1,664 | 0.025 | <0.001 | 5.07 | R | Precuneus | 7 | 28 | -46 | 46 |
| 6 | 896 | 0.020 | <0.001 | 4.24 | R | Supramarginal gyrus | 40 | 54 | -36 | 36 |
|  |  | 0.017 | <0.001 | 3.78 | R | Inferior parietal lobule | 40 | 44 | -34 | 40 |

The most significant peaks of activity were located in the left postcentral gyrus (cluster 1; TAL coordinates: -34, -30, 50, BA3), the left medial frontal gyrus (cluster 2; TAL coordinates: -6, -12, 54, BA6), the right precuneus (cluster 3; TAL coordinates: 12, -68, 48, BA7), the right middle frontal gyrus (cluster 4; TAL coordinates: 22, -8, 56, BA6), the right precuneus (cluster 5; TAL coordinates: 28, -46, 46, BA7), and the right supramarginal gyrus (cluster 6; TAL coordinates: 54, -36, 36, BA40). Cluster 1 (21,288 mm^3) showed twelve activation peaks in the left hemisphere; it extended across the postcentral gyrus (32.5% of experiments), the inferior parietal lobule (18.6%), the precuneus (15.2%), the precentral gyrus (14.6%), the superior parietal lobule (12.3%), and it slightly spread to the middle frontal gyrus (3.5%), the sub-gyral (1.7%) and the insula (1%). Cluster 2 (3,536 mm^3) consisted of three activation peaks, and it spanned both the left and right hemispheres (73.5% and 26.5% of experiments, respectively); more precisely, it was located in the medial frontal gyrus (73.5%), the cingulate gyrus (25.2%), and the paracentral lobule (1.3%). Cluster 3 (2,056 mm^3) had two activation peaks in the right hemisphere; the cluster was located primarily in BA7, and particularly, it spanned the precuneus (69.3%), and the superior parietal lobule (30.7%). Cluster 4 (1,744 mm^3) consisted of three activation peaks in the right hemisphere; it was located in the middle frontal gyrus (54.6%), the precentral gyrus (16.7%), the superior frontal gyrus (14.8%), the sub-gyral (12%), and the medial frontal gyrus (1.9%). Cluster 5 (1,664 mm^3) had one activation peak in the right BA7; in particular, it spanned the precuneus (65.3%) and the superior parietal lobule (34.7%). Cluster 6 (896 mm^3) consisted of two activation peaks in the right hemisphere; it was primarily located in the inferior parietal lobule (84.8%), and it spread slightly to the supramarginal gyrus (8.7%), and the postcentral gyrus (6.5%).

### 3.3 Contrast: Reach and Grasp Vision > No vision

The contrast meta-analysis (Reach and Grasp Vision > Reach and Grasp No vision) pooled data from a total of 528 foci, extracted from an overall group of 53 experiments and 774 participants (Reach and Grasp Vision: 168 foci, 20 experiments, 298 subjects; Reaching and Grasp No vision: 360 foci, 33 experiments, 476 subjects). The analysis revealed five significant ALE clusters. The results are represented in Figure 1 (Panel c); for more details, see Table 3.

**Table 3.** Results of the contrast analysis (Reach and Grasp Vision > No vision). TAL: Talairach; Hemi: hemisphere; BA: Brodmann Area; Cluster size: size of clusters in mm^3.

| Cluster | Cluster size | p value | z-score | Hemi | Anatomical Labelling | BA | TAL coordinates |  |  |
| --- | --- | --- | --- | --- | --- | --- | --- | --- | --- |
|  |  |  |  |  |  |  | x | y | z |
| 1 | 648 | 0.002 | 2.83 | L | Precentral gyrus | 4 | -31 | -28 | 55 |
| 2 | 576 | 0.014 | 2.19 | R | Precentral gyrus | 44 | 49 | 2 | 12 |
|  |  | 0.016 | 2.16 | R | Precentral gyrus | 44 | 47.8 | 6.5 | 11.8 |
|  |  | 0.016 | 2.13 | R | Inferior frontal gyrus | 44 | 48 | 6 | 18 |
| 3 | 344 | 0.010 | 2.31 | R | Cerebellum (anterior lobe; culmen) | - | 22 | -48 | -22 |
| 4 | 128 | 0.033 | 1.85 | R | Medial frontal gyrus | 6 | 4 | -14 | 54 |
| 5 | 112 | 0.030 | 1.88 | L | Inferior temporal gyrus | - | -48 | -70 | 2 |

The most significant peaks of activation were found in the left precentral gyrus (cluster 1; TAL coordinates: -31, -28, 55, BA4), the right precentral gyrus (cluster 2; TAL coordinates: 49, 2, 12, BA44), the culmen of the right cerebellum (cluster 3; TAL coordinates: 22, -48, -22), the right medial frontal gyrus (cluster 4; TAL coordinates: 4, -14, 54, BA6), and the left inferior temporal gyrus (cluster 5; TAL coordinates: -48, - 70, 2). Cluster 1 (648 mm^3) was found with one peak in the left hemisphere, and it was located in the precentral gyrus (BA4; 58%), and the postcentral gyrus (BA3; 42%). Cluster 2 (576 mm^3) consisted of three peaks of activation in the right hemisphere, and it was mainly located in the precentral gyrus (BA44), and the inferior frontal gyrus (BA44). Cluster 3 (344 mm^3) showed one activation peak in the right cerebellar hemisphere; it was located in the cerebellar culmen (88.4%), and slightly extended into the cerebellar dentate nucleus (11.6%). Cluster 4 (128 mm^3) revealed one activation peak in the right hemisphere, and it was located in the medial frontal gyrus (BA6; 100%). Cluster 5 (112 mm^3) showed one activation peak in the left hemisphere; it was located in the inferior temporal gyrus (61.5%), the middle occipital gyrus (30.8%), and the middle temporal gyrus (7.7%).

### 3.4 Meta-analytical map of Grasp

The Grasp ALE meta-analysis included a total of 563 subjects, and 294 foci extracted from 37 eligible experiments. Results showed five significant clusters (Figure 2, Panel a; Table 4).

**Table 4.** Results of the single dataset meta-analysis on Grasp. TAL: Talairach; Hemi: hemisphere; BA: Brodmann Area; Cluster size: size of clusters in mm^3.

| Cluster | Cluster size | ALE value | p value | z-score | Hemi | Anatomical Labeling | BA | TAL coordinates |  |  |
| --- | --- | --- | --- | --- | --- | --- | --- | --- | --- | --- |
|  |  |  |  |  |  |  |  | x | y | z |
| 1 | 12,528 | 0.051 | <0.001 | 8.35 | L | Postcentral gyrus | 3 | -34 | -30 | 56 |
|  |  | 0.038 | <0.001 | 6.89 | L | Postcentral gyrus | 2 | -42 | -28 | 50 |
|  |  | 0.038 | <0.001 | 6.78 | L | Inferior parietal lobule | 40 | -36 | -40 | 54 |
|  |  | 0.023 | <0.001 | 4.79 | L | Inferior parietal lobule | 40 | -40 | -36 | 36 |
|  |  | 0.017 | <0.001 | 3.87 | L | Precentral gyrus | 6 | -18 | -18 | 66 |
|  |  | 0.015 | <0.001 | 3.65 | L | Superior parietal lobule | 7 | -30 | -56 | 54 |
| 2 | 2,200 | 0.034 | <0.001 | 6.37 | R | Cerebellum (anterior lobe; dentate) | - | 16 | -50 | -20 |
| 3 | 2,024 | 0.025 | <0.001 | 5.14 | L | Medial frontal gyrus | 6 | 0 | -10 | 52 |
| 4 | 1,352 | 0.021 | <0.001 | 4.53 | R | Precentral gyrus | 3 | 36 | -26 | 48 |
|  |  | 0.021 | <0.001 | 4.50 | R | Postcentral gyrus | 3 | 42 | -22 | 48 |
| 5 | 1,104 | 0.024 | <0.001 | 4.96 | L | Inferior temporal gyrus | 37 | -44 | -66 | 2 |
|  |  | 0.015 | <0.001 | 3.65 | L | Fusiform gyrus | 19 | -42 | -70 | -10 |

The most significant peaks of activity were located in the left postcentral gyrus (cluster 1; TAL coordinates: -34, -30, 56, BA3), the right cerebellar dentate (cluster 2; TAL coordinates: 16, -50, -20), the left medial frontal gyrus (cluster 3; TAL coordinates: 0, -10, 52, BA6), the right precentral gyrus (cluster 4; TAL coordinates: 36, -26, 48, BA3), and the left inferior temporal gyrus (cluster 5; TAL coordinates: -44, -66, 2, BA37). Cluster 1 (12,528 mm^3) showed six significant activation peaks in the left hemisphere; it was located in the postcentral gyrus (46.7% of experiments), the inferior parietal lobule (27.5%), the precentral gyrus (22.7%), and the superior parietal lobule (1.9%). Cluster 2 (2,200 mm^3) consisted of one peak of significant activation in the right cerebellar hemisphere; the cluster was mainly located in the anterior lobe of the cerebellum (96.7%) and only slightly spread to the posterior lobe (3.3%). Cluster 3 (2,024 mm^3) was found with one peak of activation in both the left and right hemispheres (76.9% and 23.1%, respectively); the cluster was located primarily in the medial frontal gyrus (BA6; 98.5%) and activation also spread slightly to the paracentral lobule (BA31; 1.5%). Cluster 4 (1,352 mm^3) consisted of two activation peaks in the right hemisphere; it was located in the postcentral gyrus (73.3%), the precentral gyrus (25.6%), and activation also spread slightly to the inferior parietal lobule (1.2%) Cluster 5 (1,104 mm^3) was found with two peaks in the left hemisphere; it was located in the inferior temporal gyrus (45%), the middle temporal gyrus (27.5%), the middle occipital gyrus (20%), the fusiform gyrus (5%), and the inferior occipital gyrus (2.5%).

### 3.5 Meta-analytical map of Reach

The Reach ALE meta-analysis included a total of 211 subjects, and 234 foci extracted from 16 eligible experiments. Results showed six significant clusters (Figure 2, Panel b; Table 5).

**Table 5.**
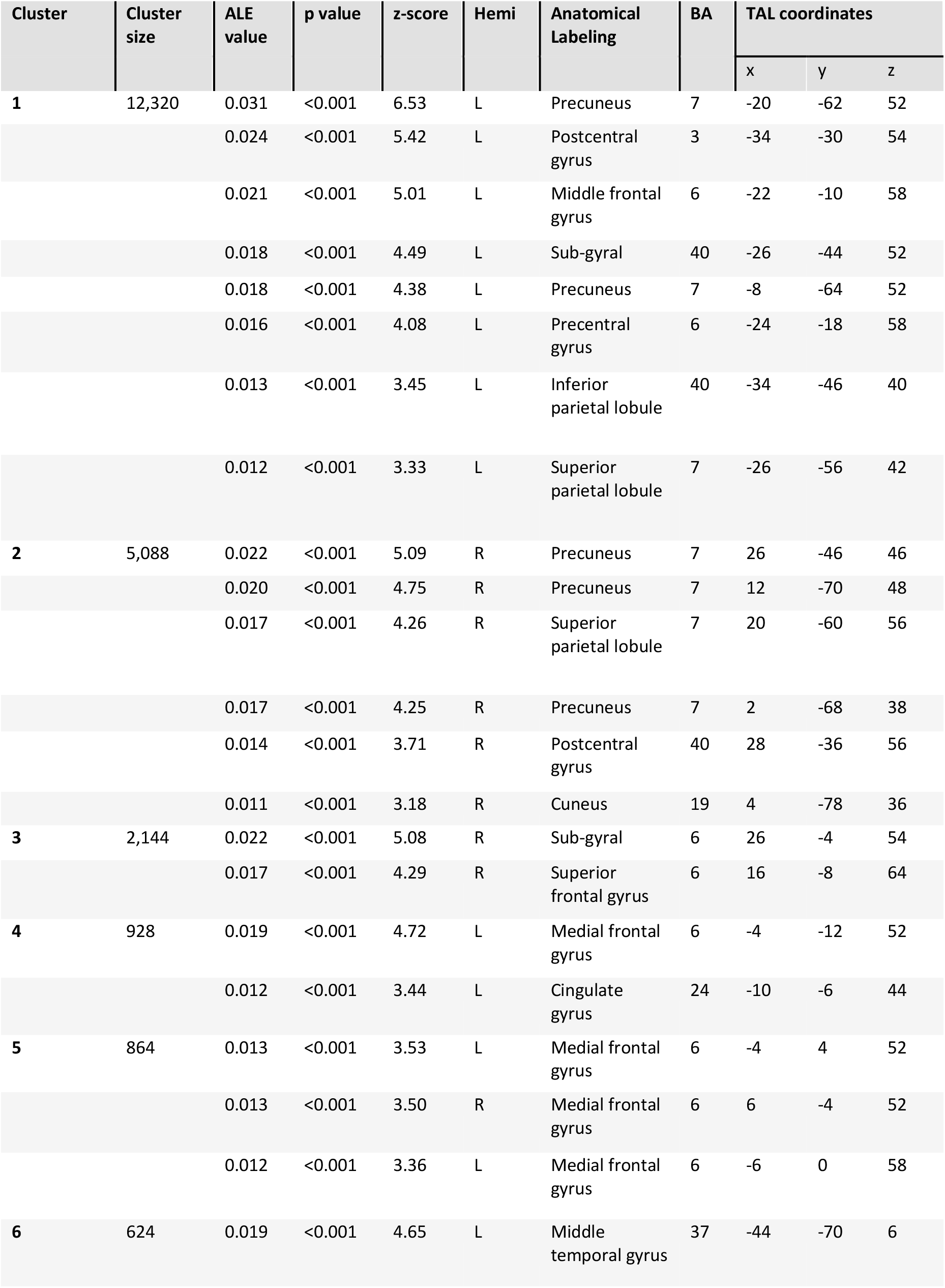
Results of the single dataset meta-analysis on Reach. TAL: Talairach; Hemi: hemisphere; BA: Brodmann Area; Cluster size: size of clusters in mm^3.

The most significant peaks of activity were located in the left precuneus (cluster 1; TAL coordinates: -20, -62, 52, BA7), the right precuneus (cluster 2; TAL coordinates: 26, -46, 46, BA7), the right sub-gyral (cluster 3; TAL coordinates: 26, -4, 54, BA6), the left medial frontal gyrus (cluster 4; TAL coordinates: -4, -12, 52, BA6; cluster 5; TAL coordinates: -4, 4, 52, BA6), and the middle temporal gyrus (cluster 6; TAL coordinates: -44, -70, 6, BA37). Cluster 1 (12,320 mm^3) showed eight peaks of activity in the left hemisphere; it was located in the precuneus (29.1% of experiments), the precentral gyrus (19%), the postcentral gyrus (18.4%), the superior parietal lobule (18.2%), the middle frontal gyrus (10.8%), the sub-gyral (2.7%), and the inferior parietal lobule (1.7%). Cluster 2 (5,088 mm^3) consisted of six activation peaks, and it spanned primarily the right hemisphere (96.6%) and, to a lesser extent, the left hemisphere (3.4%); more precisely, it was located in the precuneus (59.9%), the superior parietal lobule (24.1%), the sub-gyral (7.6%), the postcentral gyrus (6.3%), and the cuneus (1.3%). Cluster 3 (2,144 mm^3) was found with two activation peaks in the right hemisphere, particularly in BA6; the cluster spanned the middle frontal gyrus (55.3%), the superior frontal gyrus (22%), the sub-gyral (14.6%), the precentral gyrus (5.7%), and the medial frontal gyrus (2.4%). Cluster 4 (928 mm^3) consisted of two activation peaks in the left hemisphere; it was primarily located in the medial frontal gyrus (BA6; 82.9%), and the cingulate gyrus (BA24, BA31; 17.1%). Cluster 5 (864 mm^3) was found with three activation peaks in both the right and left hemispheres (50.9% and 49.1%, respectively). The latter cluster was located mainly in the medial frontal gyrus (79.2%), and activity also spread slightly to the superior frontal gyrus (13.2%), and the cingulate gyrus (7.5%). Cluster 6 (624 mm^3) showed one activation peak in the left hemisphere; it was located in the middle occipital gyrus (58.8%), the middle temporal gyrus (32.4%), and the inferior temporal gyrus (8.8%).

### 3.6 Contrast: Grasp > Reach

The contrast ALE meta-analysis (Grasp > Reach) included a total of 774 participants and 528 foci, extracted from an overall group of 53 experiments (Grasp: 294 foci, 37 experiments, 563 subjects; Reach: 234 foci, 16 experiments, 211 subjects). The analysis revealed two significant ALE clusters for activation. The results are represented in Figure 2, Panel c; for more details, see Table 6.

**Table 6.** Results of the contrast analysis (Grasp > Reach). TAL: Talairach; Hemi: hemisphere; BA: Brodmann Area; Cluster size: size of clusters in mm^3.

| Cluster | Cluster size | p value | z-score | Hemi | Anatomical Labeling | BA | TAL coordinates |  |  |
| --- | --- | --- | --- | --- | --- | --- | --- | --- | --- |
|  |  |  |  |  |  |  | x | y | z |
| 1 | 1,488 | 0.010 | 2.34 | R | Cerebellum (anterior lobe; dentate) | - | 16.8 | -51.2 | -24.8 |
|  |  | 0.015 | 2.16 | R | Cerebellum (anterior lobe) | - | 20 | -48 | -24 |
|  |  | 0.017 | 2.13 | R | Cerebellum (anterior lobe; culmen) | - | 24 | -53 | -22 |
|  |  | 0.017 | 2.13 | R | Cerebellum (anterior lobe; culmen) | - | 28 | -54 | -18 |
|  |  | 0.017 | 2.12 | R | Cerebellum (anterior lobe; culmen) | - | 22.2 | -50.9 | -16.7 |
| 2 | 184 | 0.033 | 1.85 | R | Postcentral gyrus | 3 | 44 | -24 | 52 |
|  |  | 0.037 | 1.78 | R | Postcentral gyrus | 3 | 40 | -20 | 48 |

The most significant peaks of activation were found in the right cerebellar dentate nucleus (cluster 1; TAL coordinates: 16.8, -51.2, -24.8), and in the right postcentral gyrus (cluster 2; TAL coordinates: 44, -24, 52, BA3). Cluster 1 (1,488 mm^3) was found with five peaks in the right cerebellum; this cluster was located in the anterior lobe (97.8%) and activation spread slightly into the posterior lobe (2.2%) of the cerebellum. Cluster 2 (184 mm^3) consisted of two peaks of activation in the right hemisphere, and it was primarily located in the parietal lobe, more specifically in the postcentral gyrus (BA3; 100%).

## 4. General discussion

In the present study we conducted a coordinate-based meta-analysis to investigate the brain areas consistently recruited during hand reaching and grasping with and without online visual feedback, with a focus on ventral stream and early visual areas. As for the dorsal stream, we found that it was consistently involved in grasping as well as reaching actions, regardless of the availability of visual information. This is in line with the dual-stream theory, postulating that the dorsal stream is involved in the execution of reach and grasp actions. As for the ventral stream, we found that it was involved in actions executed with but not without online visual feedback. Specifically, the temporal-occipital cortex showed higher activation likelihood for the Vision than No Vision condition. In addition, the ventral stream showed comparable activation likelihood for grasping and reaching actions. Below, we discuss the main findings of the present meta-analysis and the potential functional role of ventral stream areas in action guidance and execution.

### 4.1 Processing of visual information during hand actions

The two single-dataset meta-analyses on actions with and without vision, and the contrast analysis between Vision and No Vision enabled us to examine the role of vision in occipital-temporal areas during action execution (aim 1).

The meta-analytical map on hand reaching and grasping with vision (Figure 1, Panel a) revealed consistent activation across the studies in frontal, parietal, and right cerebellar regions, as well as in the occipito-temporal cortex. These findings are expected given the known role of the dorsal stream in action and the ventral stream in perception. Indeed, the inferior temporal gyrus (ITG) and the lateral occipital cortices are known to be involved in visual perception and recognition of object categories and body parts, including the hand (Bracci et al., 2010; Herath et al., 2001; Ishai et al., 1999; Malach et al., 1995; for reviews see Grill-Spector, 2003; Grill-Spector & Weiner, 2014). Similarly, the posterior areas of the middle temporal gyrus (MTG) process category and motion-related information (Chao et al., 1999). These results are consistent with the fact that in the selected Vision conditions participants viewed the target and their moving hand while approaching the target.

As for actions performed without online visual feedback, the meta-analytical map showed significant activation in six cortical clusters covering both the left and right hemispheres; specifically, they included parietal areas, such as bilateral inferior parietal lobule (IPL), precuneus, superior parietal lobule (SPL), sub-gyral, and left postcentral gyrus (PoCG), and frontal areas like bilateral precentral gyrus (PreCG), middle frontal gyrus (MFG), superior frontal gyrus (SFG), left insula. Importantly, no cluster was found in the ventral visual stream (Figure 1, Panel b). Therefore, despite some evidence supporting the involvement of the ventral stream in hand actions performed in the dark after a delay (Monaco et al., 2017; Singhal et al., 2013), there is no consistency in support of the recruitment of ventral visual stream areas during the control and execution of skilled actions in the absence of visual input. In line with this result, the contrast analysis did not reveal any cluster in the temporal-occipital cortex (Figure 1, Panel c). Interestingly, the contrast did reveal clusters of activation spanning the left PreCG and PoCG, two clusters in the right hemisphere located mainly in frontal areas, such as PreCG, inferior and middle frontal gyrus (IFG and MFG, respectively), and an additional cluster in the anterior lobe of the right cerebellar hemisphere. Therefore, these areas are more engaged when visual information is available as opposed to when it is not. The higher consistency in brain activation for Vision than No Vision in motor-related areas supports the strong involvement of the dorsal stream and premotor areas in the online visual guidance of actions. A seminal neurophysiology study in primates by Graziano and colleagues (1997) demonstrated the presence of bimodal visual and tactile neurons in macaques’ ventral premotor cortex, typically known to be involved in motor control. Our results suggest that bimodal neurons might also be present in humans’ premotor cortex, which is recruited by vision of a target to be acted upon that requires the processing of affordances and spatial information for accurate action performance.

We found no activation cluster in the early visual cortex for Vision and No Vision condition. The lack of activation clusters in the early visual cortex in the Vision condition can be explained by the fact that some of the studies included in the Vision condition used a contrast of Grasp > Look. Therefore, the visual processing of the object was subtracted from the activity in the early visual cortex allowing the observation of activation areas involved in the execution of action when vision is not prevented. We included these studies in the Vision condition because the contrast did not subtract the visual processing of the grasping hand however, the inclusion of contrasts subtracting some activity related to visual processing might have reduced the sensitivity to detect activation in early visual areas. Also, while univariate analysis might lack the sensitivity to reveal activation in ventral stream areas and early visual cortex under lack of vision, recent multivoxel pattern analysis studies have shown different representations for grasping and reaching action planning with and without visual information in ventral stream and early visual areas (Monaco et al., 2019, 2021). This difference in results indicates that univariate and multivariate analysis provide complementary and not necessarily equivalent information, with MVPA being more sensitive to distributed representation of information of content, and univariate analysis showing more sensitivity to the overall engagement in a task (Coutanche, 2013; Davis et al., 2014; Jimura & Poldrack, 2012).

Overall, these results indicate the consistent involvement of the parietal and frontal cortex in the execution of grasping and reaching actions regardless of the availability of visual information, while the temporal-occipital cortex is recruited only when vision is available. Notably, the same frontal and parietal areas have been recently shown to process magnitude representations (Cona et al., 2021). Consistently, grasping and reaching actions require the processing of space-related magnitude information, such as the location of the target in space for reaching, as well as its size for grasping.

### 4.2 Processing of grasping and reaching

The second aim of the current study was to assess whether ventral and dorsal stream areas are differentially recruited during the execution of reaching and grasping (aim 2).

The meta-analytical map of hand grasping revealed consistent activation clusters in the frontal and parietal cortex, as well as in the anterior lobe of the right cerebellar hemisphere (Figure 2, Panel a). Moreover, a significant cluster of activation was located in the left inferior temporal cortex, more precisely in the ITG, and the fusiform gyrus. The inferior temporal cortex, including the ITG, plays a well-known role in the visual perception and recognition of objects, scenes, and hands (Bracci et al., 2010; Grill-Spector & Weiner, 2014; Kanwisher, 2010; Malach et al., 1995) and it has been shown that the fusiform gyrus might store semantic information about the object shape (Chao et al., 1999; see also Sakreida et al., 2016). Therefore, neural activation in these areas might underpin the visual processing of object properties and semantic representation.

Similarly, the meta-analytical map of hand reaching revealed five cortical clusters covering both hemispheres (Figure 2, Panel b). Specifically, a network of frontal, parietal, and temporal-occipital regions was consistently activated across the literature. Moreover, the network for reaching appeared more bilateral than the one for grasping, in line with previous findings showing a lateralization of the activation for grasping movements (Blangero et al., 2009; Ranzini et al., 2022). Concerning the involvement of dorsal stream areas, slight differences can be seen in the areas included in the grasping and reaching networks, which might probably be associated with different types of processes involved in the different types of actions as recently discussed by Ranzini et al. (2022).

Overall, these meta-analyses revealed that both dorsal and ventral stream areas are recruited during hand reaching and grasping across the two visual conditions.

### 4.3 Strengths and limitations of the study

The present meta-analysis has important strengths, as it is the first work to define the consistency across neuroimaging studies on hand actions with and without online visual information over the last two decades. In addition, this study provides a further confirmation of the involvement of dorsal stream areas in the control of skilled actions (Culham et al., 2006; Gallivan & Culham, 2015; for additional meta-analytical results, see also Ranzini et al., 2022). Furthermore, we provide an overview of the dorsal and ventral stream areas recruited while reaching out and grasping objects, with and without online visual feedback. This aspect is of particular importance. Indeed, despite the fact that there is consensus in the literature on the involvement of the dorsal stream in the guidance of reaching and grasping, there is compelling evidence suggesting that also the ventral stream might play a role (e.g., Cohen et al., 2009; Milner & Goodale, 2008; Monaco et al., 2017; Singhal et al., 2013), especially in recent neuroimaging studies that have employed multivariate analysis which allows identifying representations rather than extent of activation (Gallivan et al., 2013, 2019; Gutteling et al., 2015; Monaco et al., 2019, 2021). Therefore, our meta-analysis attempted to clarify the debate on the potential involvement of the occipito-temporal cortex in the guidance of skilled actions as investigated with univariate analysis.

There are also some limitations to our work. One of them consists in the fact that half of the experiments categorized as ‘Grasping with vision’ used a contrast (Grasp > Look/Reach/Point/Touch/Different grasp) which might have removed visual processing elicited by the vision of the moving hand and/or the object, as well as areas involved in processing arm movement (Grasp > Reach). This issue might have hampered the possibility to find consistent activation in brain areas associated with visual processing during action execution in presence of vision. As a consequence, it might have likely hindered the contrast analysis between the two vision conditions (i.e., Reach and Grasp Vision > No vision) which did not show any significant cluster of activation associated with visual processing in the early visual cortex.

## 5. Conclusion

To the best of our knowledge, this coordinate-based meta-analysis is the first attempt to investigate spatial convergence across the available literature in relation to the involvement of ventral and dorsal visual streams in reaching and grasping actions performed with and/or without online visual feedback. Our findings reconcile the existing neuroimaging literature on actions that employed standard univariate analysis, by emphasizing the complementary role of more recent techniques, such as multivoxel pattern analysis, to the current knowledge on cortical areas involved in hand movements.

## Acknowledgements

The authors would like to thank Marco Bedini for useful discussion.

This research did not receive any specific grant from funding agencies in the public, commercial, or not-for-profit sectors.

## Author Contribution

**Samantha Sartin**: Data curation; Formal analysis; Methodology; Project administration; Resources; Software; Visualization; Roles/Writing - original draft. **Mariagrazia Ranzini**: Investigation; Data curation; Writing - review & editing. **Cristina Scarpazza**: Writing - review & editing. **Simona Monaco**: Conceptualization; Methodology; Project administration; Resources; Supervision; Validation; Writing - review & editing.

## Annexes

Annex A: PRISMA Flow diagram for studies on reaching and grasping (adapted from Ranzini et al., 2022).

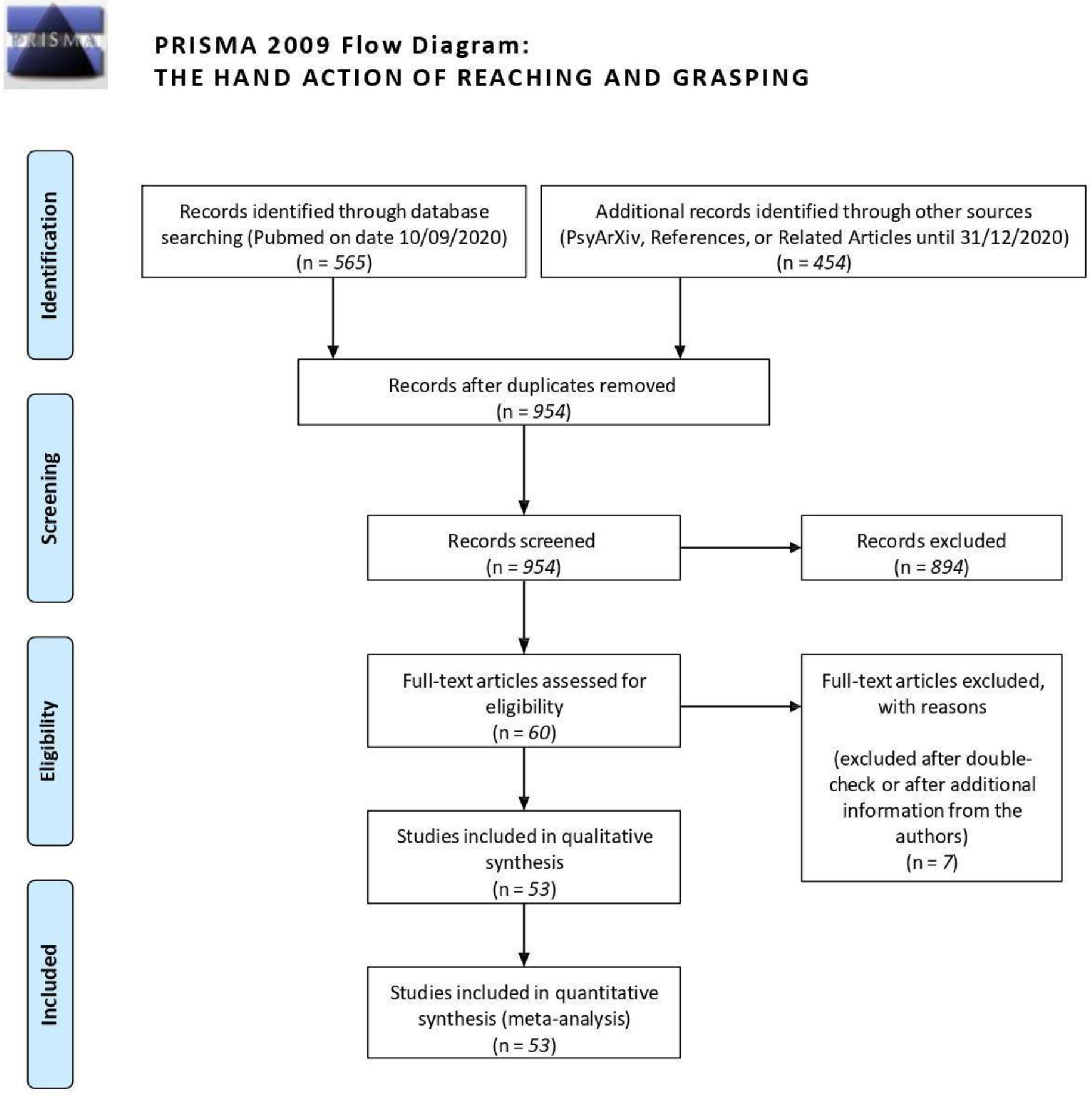

Annex B: Table of the list of included studies and additional details about participants, the task employed, contrast used, category, and number of activation foci (adapted from Ranzini et al., 2021).

| Study N° | Study (First author, publication year) | N, Sex, Handedness | Age | Technique | Contrast | Category | Details | N Foci |
| --- | --- | --- | --- | --- | --- | --- | --- | --- |
| 1 | Styrkowiec, 2019 | 21, 11 F, all RH | 23 | fMRI | grasp tool > control object | Grasp No vision | Execution w/o vision; real object tools and non-tools | 19 |
| 2 | Chen, 2018 | 12, 8 F, all RH | 33 | fMRI | Response: different cue > same cue | Reach No vision | Reach-to-touch-with index finger | 9 |
| 3 | Gorbet, 2018 | 20, 20 F, all RH | 25 | fMRI | Reach > dot fixation | Reach Vision | Visuo-motor combination | 15 |
| 4 | Gertz, 2015 | 19, 11 F, all RH | 25 | fMRI | Reach-Pro > Reach underspecified | Reach No vision | Pro and Anti reach task in specified (with earlier instruction) or underspecified (with later instruction) condition | 7 |
| 5 | Renzi, 2013 | 9, 5 F, all RH | 24 | fMRI | Far > Near in visual feedback | Grasp No vision | Near or far whole or precision grasp with or without visual feedback | 4 |
| 6 | Rossit, 2013 | 10, 7 F, all RH | 27 | fMRI | Grasp in Lower > Upper visual field in absence of vision | Grasp No vision | Grasp or look 3D objects central objects with focus on the lower vs. upper visual field | 3 |
| 7 | Bernier, 2012 | 18, 7 F, all RH | 26 | fMRI | Uncued > Cued (right hand) | Reach No vision | Short cued or uncued reach with either left or right hand to left or right targets | 9 |
| 8 | Glover, 2012 | 21, all RH |  | fMRI | Immediate execution > Observation | Grasp Vision | Four conditions: Observation, Imagination (planning), Immediate execution (Control), Planning + Control | 11 |
| 9 | Fabbri, 2012 | 13, 8 F, all RH | 27 | fMRI | changed > adapted test trials | Reach No vision | Adaptation reach direction or amplitude | 18 |
| 10 | Gallivan, 2011 | 13, 7 F, all RH | 28 | fMRI | Reach & Grasp > Look (right handers) | Grasp No vision | Reach, Grasp or Look within near or far space in right or left handers | 1 |
| 11 | Fiehler, 2011 | 21, 15 F, all RH | 24 | fMRI | Immediate grasp > baseline | Grasp Vision | Reach grasp without visual feedback, immediately or after a delay | 7 |
| 12 | Fabbri, 2010 | Exp 1: 14, 6 F, 13 RH | 28 | fMRI | changed in direction or type of motor act > adapted test trials | Reach No vision | Press or whole grasp (both with reach varying in direction) | 8 |
| 13 | Himmelbach, 2009 | 16, 8 F, all RH | 32 | fMRI | Immediate reach > delayed reach | Reach Vision | Reach with left or right hand (healthy control participants) | 11 |
| 14 | Verhagen, 2008 | 19, 0 F, all RH | 22 | fMRI | Grasp in binocular & monocular vision > rest | Grasp Vision | Monocular or binocular visually guided precision grip | 10 |
| 15 | Begliomini, 2007 | 16, all RH | 25 | fMRI | Precision grip > power grasp | Grasp Vision | Reach to grasp small or large objects with precision or power grip | 1 |
| 16 | Valyear, 2019 | 17, 0 F, all RH | 45 | fMRI | Grasp > Touch | Grasp Vision | Brain activity of the healthy control group | 2 |
| 17 | Filimon, 2007 | 14, all RH | 34 | fMRI | Reach Execution > Object Viewing | Reach No vision | Reach execution, reach observation, reach imagination | 35 |
| 18 | Milner, 2007 | 17 |  | fMRI | Complex > Simple grasp | Grasp No vision | Squeezing a soft ball (simple) or balancing a weighted flexible ruler (complex), or rest (precision grip only) | 2 |
| 19 | Begliomini, 2007 | 12, 8 F, all RH | 25 | fMRI | Precision grip > Whole hand grasp | Grasp Vision | Precision grip or whole hand grasp | 1 |
| 20 | Ehrsson, 2007 | 6, 0 F | 26 | fMRI | Loading > Rest | Grasp No vision | Precision grip during increasing (loading) or decreasing (unloading) in weight | 3 |
| 21 | Prado, 2005 | 12, 8 F, all RH | 23 | fMRI | Reach w/o saccade > No reach | Reach No vision | Reaching with saccade, reaching w/o saccade (VT/NSe), reaching with invisible object | 13 |
| 22 | Frey, 2005 | 14, 5 F, all RH | 24 | fMRI | Grasp > Point | Grasp Vision | Exp.2: Precision grasp | 2 |
| 23 | Kuhtz-Buschbeck, 2001 | 8, 0 F, all RH | 29 | fMRI | Gentle > Firm grasp | Grasp No vision | Precision grip with gentle, normal or firm grip force | 4 |
| 24 | Cavina-Pratesi, 2018 | 11, 4 F, all RH | 31 | fMRI | all Reach / Point > Passive viewing | Reach No vision | Reach, point, precision or coarse grip with two or more digits | 11 |
| 25 | Monaco, 2015 | 11, 5 F, all RH | 32 | fMRI | Adaptation to both size and location | Grasp Vision | Grasp specific adaptation to object size and location | 5 |
| 26 | Monaco, 2014 | 13, 8 F, all RH | 31 | fMRI | Adaptation to Grasp > Adaptation to View | Grasp Vision | Grasp specific adaptation to grasp type or object size | 3 |
| 27 | Monaco, 2011 | 11, 3 F, all RH | 33 | fMRI | Adaptation in Grasp > Adaptation in Reach | Grasp Vision | Adaptation to object orientation during grasp, reach or look | 5 |
| 28 | Cavina-Pratesi, 2010 | Exp.1: 10, all RH | 29 | fMRI | Grasp > Touch (Exp.1) | Grasp Vision | Grasp, touch and look to near or far objects | 11 |
| 29 | Cappadocia, 2017 | 12, 9 F, all RH | 28 | fMRI | Pro & Anti Reach > Color detection (execution phase) | Reach No vision | Pro and anti reach task | 17 |
| 30 | Pellijeff, 2006 | 13, 8 F, 12 RH | 25 | fMRI | First reach > Later Reaches | Reach No vision | Chin pointing with the index finger (reach, exp.1) | 6 |
| 31 | Króliczak, 2007 | 10, 5 F, all RH | 27 | fMRI | Grasp > Reach | Grasp No vision | precision grasp, reach, pantomime grasp, pantomime reach | 12 |
| 32 | Inoue, 1998 | 9, all RH | 23 | PET | Reach without feedback > Hold (control task) | Reach No vision | Reach without feedback | 30 |
| 33 | Desmurget, 2001 | 7, 1 F, all RH | 25 | PET | Reach > Look (stationary condition) | Reach No vision | Reach or look to stationary or jumping targets | 15 |
| 34 | Kertzman, 1997 | 6, 3 F, all RH | 33 | PET | Reach with right hand > Look | Reach Vision | Reach with index finger; right or left hand on right or left visual field | 14 |
| 35 | Keisker, 2009 | 14, 7 F, all RH | 27 | fMRI | Grasp > Rest | Grasp Vision | Power grip with three forces | 15 |
| 36 | Keisker, 2010 | 14, 7 F, all RH | 27 | fMRI | Grasp static & dynamic > No force | Grasp Vision | Power grip in static or dynamic condition | 12 |
| 37 | Ward, 2003 | 26, 9 F, all RH | 47 | fMRI | Effect of grip force | Grasp No vision | Power grip with left or right hand | 6 |
| 38 | Hilty, 2010 | 15, 0 F, all RH | 25 | fMRI | Succeeded grasp > baseline | Grasp Vision | Power grip with different forces and during interference | 5 |
| 39 | Kurniawan, 2010 | 17, all RH | 27 | fMRI | Grip low effort > grip high effort | Grasp No vision | Power grip or hold with high or low effort, and with or without reward | 10 |
| 40 | Talelli, 2008 | 27, all RH | 42 | fMRI | Effect of grip force | Grasp No vision | Dynamic power hand grips with the dominant right hand | 4 |
| 41 | Vaillancourt, 2003 | 10, 7 F, all RH | 27 | fMRI | Grip with visual feedback > rest, no visual feedback or visual stimulus | Grasp No vision | Precision grip with visual feedback, without visual feedback, visual stimulus only or rest | 26 |
| 42 | Spraker, 2009 | 12, 7 F, all RH | 27 | fMRI | Grip > relaxation | Grasp No vision | Precision grip or relaxation phase | 1 |
| 43 | Neely, 2013 | 17, 8 F, 16 RH | 27 | fMRI | Dynamic grip > static grip | Grasp No vision | Static or dynamic precision grip | 13 |
| 44 | Holmström, 2011 | 16, 0 F, all RH | 32 | fMRI | High grip force > low grip force | Grasp Vision | Precision grip with high or low force, and high or low instability | 12 |
| 45 | Saiote, 2016 | 31, 15 F, all RH | 32 | fMRI | Action execution > imagination | Grasp No vision | Squeezing a ball or imagine doing it | 15 |
| 46 | Turella, 2009 | 16, all RH | 28 | fMRI | Hand action > Look | Grasp Vision | Reach and grasp with precision grip a small ball, or fix the ball | 4 |
| 47 | Gatti, 2017 | 24, all RH | 23 | fMRI | Grasp > Simple movement | Grasp No vision | Simple (palm movement), Complex (finger movement), Finalistic (Grasp) hand actions | 6 |
| 48 | Bernier, 2017 | 15, all RH | 23 | fMRI | Reach > Finger movement | Reach No vision | Reach a target with the index finger or finger movement in response to target position (no point) | 17 |
| 49 | Begliomini, 2015 | 16, all RH | 25 | fMRI | Effect of Grasp | Grasp Vision | Precision grasp with right or left hand | 22 |
| 50 | Fabbri, 2014 | 15 | 37 | fMRI | Execution > baseline | Grasp No vision | Precision or power grasp, or touch | 10 |
| 51 | Ehrsson, 2003 | 6, all RH | 27 | fMRI | Grip & lift > lift only | Grasp No vision | Grip and try to lift, grip, or try to lift w/o grip | 7 |
| 52 | Marangon, 2016 | 10, 4 F, all RH | 28 | fMRI | Grasp > Reach | Grasp No vision | Reach or reach and grasp unfamiliar simple or complex objects after exploration | 4 |
| 53 | Monaco, 2017 | 18, 7 F, all RH | 29 | fMRI | Delayed Grasping > Delayed Reaching (execution phase) | Grasp No vision | Haptic or visual grasp and/or reach task | 16 |

